# Fast-timescale hippocampal processes bridge between slowly unfurling neocortical states during memory search

**DOI:** 10.1101/2025.02.11.637471

**Authors:** Sebastian Michelmann, Patricia Dugan, Werner Doyle, Daniel Friedman, Lucia Melloni, Camilla K. Strauss, Sasha Devore, Adeen Flinker, Orrin Devinsky, Uri Hasson, Kenneth A. Norman

## Abstract

Prior behavioral work showed that event structure supports our ability to search through continuous naturalistic memories. We hypothesized that, neurally, this memory search process involves a division of labor between slowly unfurling neocortical states representing event knowledge and fast hippocampal-neocortical communication supporting retrieval of new information at transitions between events. In a sample of ten intracranial EEG patients we tracked slow neural state-patterns as patients viewed a movie and then searched their memories in a structured naturalistic interview. As patients answered questions (“after A, when does B happen next?”), state-patterns from movie-viewing were reinstated in neocortex. During the critical memory-search period, states unfurled in a forward direction, realizing a trajectory from A to B in memory. Punctate moments of state-transition during memory-search were marked by low-frequency power decreases in cortex and preceded by power decreases in hippocampus that correlated with reinstatement. Connectivity-analysis revealed information-flow from hippocampus to cortex underpinning state-transitions.

## Main text

Humans have the ability to relive the past in rich detail^1^. As you recall a cherished memory of a beach day, you can picture yourself building a sand castle during low tide and then you can picture the waves splashing over it when the tide comes in, washing it away. Intuitively, there are several similarities between such vivid memories of continuous experience and a video that we re-watch: both are records of the past that contain rich details, span long periods of time, and support sequential replay.

On a neural level, the process of memory retrieval is supported by the hippocampal formation^2, 3^. Furthermore, numerous studies have shown that remembering entails reinstating neural activity patterns that characterize the original perception in neocortical regions^4, 5^. But what neural mechanisms support the retrieval of memories of temporally extended events like a day at the beach? One possibility is that the hippocampus could act like a neural movie projector that continuously sends information back to the cortex. In this view, the neocortex assumes a passive role and does not store knowledge that could contribute to the retrieval process. In contrast, this view credits the hippocampus with holding a frame-by-frame copy of the original experience. Such video-like memories would predict a steady flow of information from hippocampus to neocortex underpinning continuous retrieval.

Empirical evidence paints a more complex picture; continuous memories are, in fact, strikingly different from videos. Memories of long narratives are organized in memory based on their structure: The continuous beach day can be broken up into so-called *events* – meaningful units of activity like “putting on sunscreen”, or “going for a swim”^6–12^. Moreover, studies that assess the time that it takes to remember an experience routinely find that (unlike a video) recall is substantially faster than the original experience and crucially that it is the event structure of the memory trace that modulates the speed at which it is replayed^13–18^. Indeed, recent behavioral data suggests that event structure allows us to dynamically access our memories throughout a continuous retrieval process, providing them with an organization akin to chapters, where it is possible to “skip ahead” to the next event^14^. Studies using functional magnetic resonance imaging (fMRI) have further shown that, during perception, high-level cortical regions break up continuous sensory input into a sequence of neural event representations. Specifically, the pattern of neural activity that characterizes a given event remains stable throughout the event and then shifts abruptly at event boundaries^19–21^. Different events of the same kind (e.g., two distinct events of swimming in the ocean) are characterized by similar event representations in cortical areas^22–24^, suggesting that these cortical event patterns may represent generalized event knowledge – an “event model”; crucially, these event models are accessed both during perception and also during recall of naturalistic narratives^19, 25^. According to theories of event segmentation, event models contain rich knowledge of what will happen next within an event, based on repeated prior experiences with that event type^7, 26^. Taken together, these points underline the importance of accounting for cortical event knowledge as well as hippocampus-driven reinstatement when explaining continuous memory retrieval.

The idea that cortex contains knowledge of the structure *within* events suggests that there is relatively less need to retrieve information from the hippocampus during an event (since cortex can support this retrieval to a larger extent), and more need for hippocampal retrieval at the boundaries between events, where there is more uncertainty about what will happen next^27–29^; at these moments, hippocampus can provide previously stored information to neocortex that is not available within the current event model^30^. In support of this view, a recent study of patients undergoing intracranial electroencephalography (iEEG) recording for clinical purposes found that when patients listened to a story for the second time, enhanced hippocampocortical information flow supported the recall of upcoming information around the time of event boundaries^31^. This prior observation of predictive recall near event boundaries during story-listening suggests that the event structure of experience may also play a crucial role when we mentally scan through continuous memories of past events. We perform this kind of memory-scanning routinely when we search through memory to answer questions about the past (e.g., in thinking about when we last had our keys); however, the fast time-scale neural mechanisms of unconstrained memory-search are poorly understood. We hypothesize that, during memory-search, continuous memories are unfurled through a complex division of labor between hippocampus – holding information about the sequence of events and unique details within those events – and neocortex – contributing its generalized event knowledge to the retrieval process^32^. The signature prediction of this view is that hippocampal contributions to a continuous retrieval process should not be uniform in time; rather, they should be especially strong at transition points between events (i.e., event boundaries), where hippocampal retrieval “feeds” neocortex information about the properties of the upcoming event. According to this view, the strength of hippocampal activation at an event transition should predict the fidelity of cortical pattern reinstatement in the subsequent event, and information transfer from hippocampus to neocortex should be a hallmark of event transitions in memory.

To test these predictions, we relied on concurrent recordings from hippocampal and neocortical channels in a rare sample of ten iEEG patients with well-preserved episodic memory function; this allowed us to measure neural activity with high spatial and temporal precision. In our study, patients watched a shortened movie (Gravity, Cuaron 2013) in two parts. After each half, they engaged in a naturalistic interview with the experimenter (Figure 1). The experimenter first described a scene in the movie (e.g., “remember the flames flying into the hallway”), confirming with the patient that they remembered that scene. Subsequently, the patient was asked about a later scene in the movie (e.g., “… when is the next time we see fire?”). The time between the end of the question and the beginning of the patient’s answer marked a memory-scanning period, during which the patient searched their memory for an answer without any external constraints. Importantly, the wording of the questions in the interview (“when is the next time…”) was carefully chosen to induce forward scanning through the event, and prior behavioral work using this paradigm indicates that this was the case^14^. This period of silent memory-scanning can last for several seconds before participants report their answer. Our focus was on “looking inside the black box” during this period to shed light on how participants mentally advance from one event memory to another during naturalistic memory search. Notably, the absence of stimuli during this period sets our work apart from studies that have explored bridging between events during stimulus perception; and the absence of responding during this memory-scanning period sets our work apart from free recall studies, where the demand to verbally describe retrieved events one-by-one during the process of memory search may distort natural retrieval dynamics.

**Figure 1:**
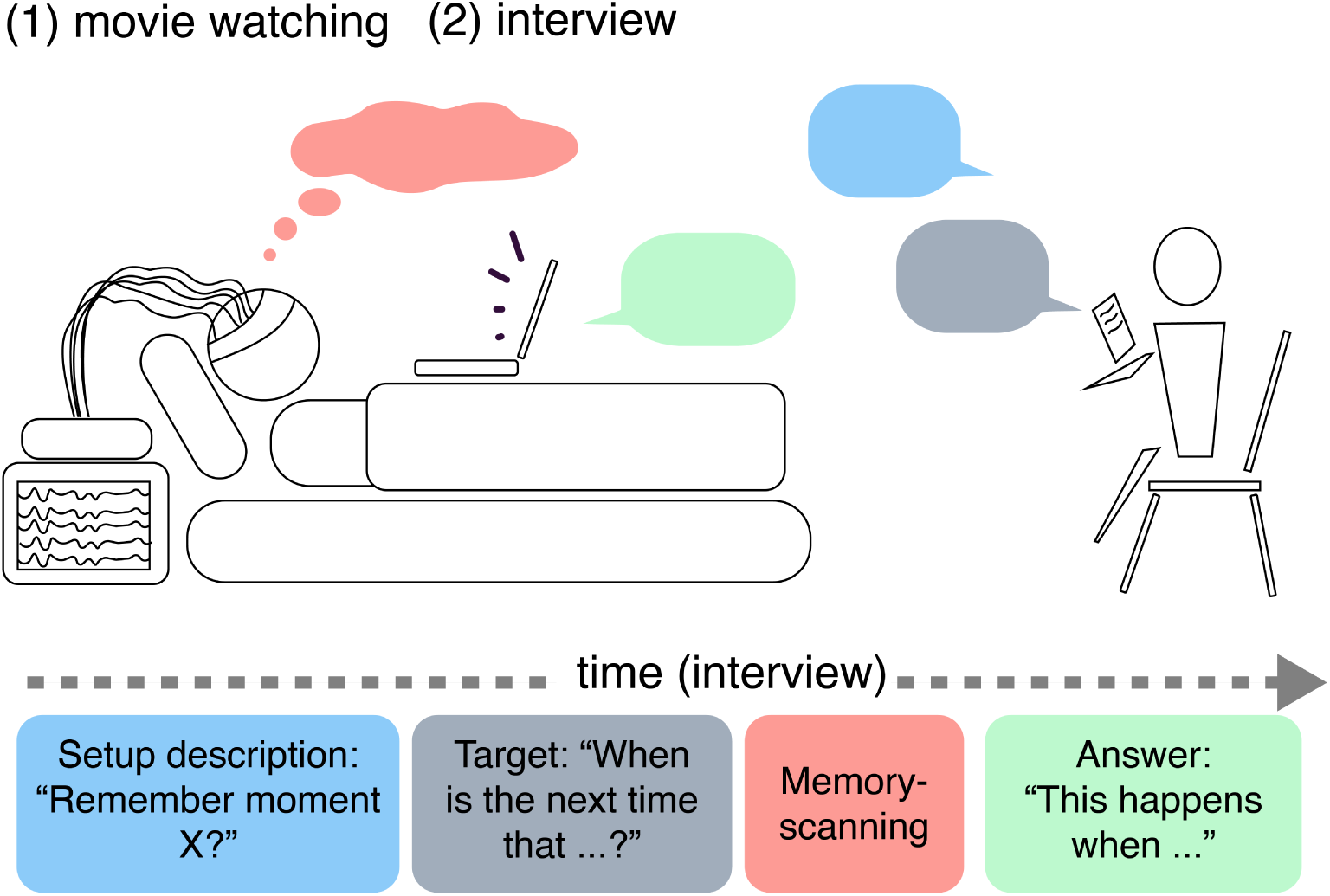
Experimental paradigm. Patients watched the two movie parts on a laptop in the hospital while undergoing intracranial electroencephalography recording for clinical purposes. After each movie, the experimenter asked naturalistic interview questions that consisted of a setup and a target. The setup described a scene in the video, the target asked about a later scene. The wording of the questions was crafted to induce forward memory scanning (see^14^ for behavioral evidence that this was indeed the case). Patients then searched their memory and answered by describing the correct scene verbally. The time between the description of the target and the first word in the patient’s answer marks the memory-scanning period. This memory-scanning period is a crucial period of interest, allowing us to study interactions between hippocampus and cortex when no external stimulus is driving perception or memory.

### Neural event segmentation

A key challenge in studying the role of events in this type of unconstrained memory-scanning was to identify when transitions between idiosyncratic event patterns occur. Unlike the manual annotations that can be derived from story stimuli during story-listening or from the transcripts of free recall^25, 31, 33, 34^, there is no external behavior to mark event transitions during memory-scanning. We addressed this challenge by identifying event boundaries in a data-driven way during movie-viewing as the moments of transition between slowly-changing neural states^19, 20^. Doing this segmentation on iEEG data involves its own set of challenges – electrophysiology is typically characterized by fast timescale dynamics, but here we sought to segment experience into longer-lasting neural events, which required us to identify slowly-changing components of the iEEG data. To accomplish this goal, we used time delay embedding, a foundational method in nonlinear time series analysis that transforms recordings into a higher-dimensional representation that reveals the system’s latent structure and enables the reconstruction of long-timescale organization from short-timescale fluctuations^35^. In brief, our procedure was composed of two steps: The first step involved concatenating lagged versions of the iEEG time series from a local neighborhood of five channels, effectively replacing spatial patterns with spatio-temporal patterns spanning a time window of 250 ms^36^. We then applied a novel dimensionality reduction to the embedded signal that (in essence) extracts principal components from its smoothed autocovariance, maximizing the temporal stability of extracted components (see Methods). Tailored dimensionality reduction techniques are a powerful tool for reducing cognitively relevant aspects of high-dimensional data to a lower-dimensional space^37^; the net effect of the steps we took here was to transform the raw iEEG data time series into components that capture spatio-temporal patterns that are stable over time spans of multiple seconds. This transformation was applied in a cross-validated way. Slow components were subsequently segmented into discrete chunks at time-points detected via Greedy-State-Boundary-Search^20^, which maximizes the difference between within-segment and between-segment similarity.

### Neural event boundaries are aligned with behavioral event segmentation

Based on prior work with fMRI^19^, we expected that this data-driven neural segmentation would align with behavioral event segmentation judgments. To validate our neural event segmentation and to confirm that the slow spatiotemporal patterns we extracted captured meaninful event information, behavioral data from 203 participants were collected via Amazon’s Mechanical Turk. In brief, participants in this norming-sample pressed a button whenever they found that one natural and meaningful unit in the movie ended and another began^7^. Observers agreed substantially on these moments (Figure 2a) and local peaks in agreement were considered behavioral event boundaries (see^14^ for a detailed description of the norming study). To measure the association between neural state-boundaries and behavior, we treated the perception of event boundaries in the behavioral norming study as a continuous variable: The proportion of participants who pressed a button within each second of the movie was taken as the strength of agreement on an event boundary (see, however, Figure 2b for depictions of discretized behavioral event boundaries together with examples of neural state boundaries). We next computed the average behavioral agreement relative to each channel’s neural state-boundaries (i.e., we locked the behavioral time-course of agreement to the time-points of state-boundaries in the neural data). We then z-scored the time-locked agreement per channel relative to a null distribution created by permuting the order of events (see Methods) and averaged these z-scores across all channels for each patient. Note that, under the null hypothesis, these time-courses of z-scores will average to zero; however, we observed a significant increase in behavioral agreement across patients that appeared *after* neural state-boundaries and peaked at 1261 ms after transitions (i.e., button presses in the norming-sample tended to follow neural state-boundaries derived from the patient data set by approximately 1.3 seconds). Agreement was significantly increased at many time-points between 446 ms and 6023 ms, *p*s *<*0.0108, controlling the false discovery rate at *q* = 0.025; cluster permutation revealed one significant cluster of increased agreement between 155ms and 8343 ms after neural state-boundaries, *p <*0.001 (Figure 2c, Supplementary Figure 3). To interpret the strength of this effect at each channel, we computed a noise ceiling from the behavioral data only. Specifically, we repeated the above analysis locking agreement to the behavioral event boundaries derived from the same data (local peaks marked by dashed lines in Figure 2a), obtaining the maximum z-score for an ideal state segmentation. We then expressed each channel’s maximum z-score within the time-window of the significant cluster as a percentage of that noise ceiling (Figure 2d, maximum value: 27.72%).

**Figure 2:**
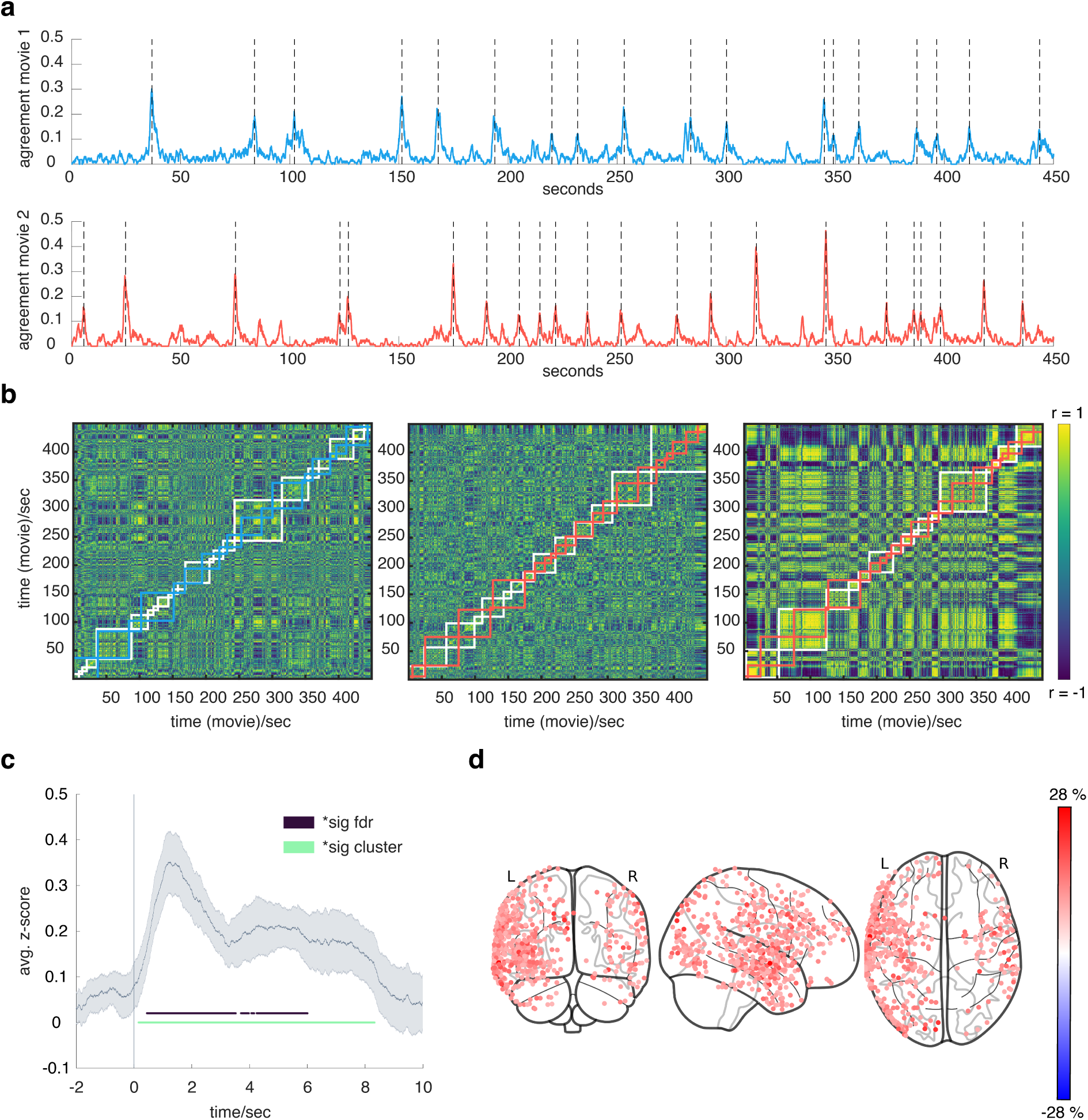
Neural state segmentation tracks shared event perception. **(a)** Agreement on event boundaries in the norming sample for the first (top; blue) and second (bottom; red) half of the movie. Colored lines indicate the proportion of participants that pressed the response button within each second; dashed vertical lines mark local peaks in agreement. **(b)** Examples of state segmentation on three channels. White lines are overlaid on time by time correlation matrices of slow components; these lines delineate neural state boundaries. Overlaid colored lines delineate event boundaries defined by the behavioral norming sample on corresponding parts of the movie (blue = first, red = second; based on dashed lines). **(c)** Grand average z-score (± SEM) of the continuous measure of agreement from (a), locked to neural state boundaries (average of averages across channels and patients; SEM across patients). Colored dots indicate points of significant increase controlling the false discovery rate (purple) and derived from a cluster permutation (green). Agreement peaks 1261 ms after neural state-boundaries. **(d)** Peak association of neural state-boundaries with behavior, expressed as percentage of the noise ceiling. The plot shows channels where the maximum z-score exceeded *Z*_0.95_ = 1.645 in the significant temporal cluster shown in green in part c.

### Neural event states are reinstated during the naturalistic interview

Our primary goal was to study neural reinstatement of event patterns during the silent memory search period. However, before doing that, we first tested whether neural states defined by segmentating the movie-viewing data were reinstated during the parts of the naturalistic interview when the experimenter or patient was talking (i.e., excluding the silent memory search period). We analyzed those questions that were unambiguously answered correctly by the patients (mean performance = 57.78%, median = 55.56%, SD = 20.49%, min = 27.78%, max = 88.89%). Other answers were highly heterogenous: They consisted of unclear descriptions, follow-up questions by the patient, long silences where no answer was provided, and descriptions of the wrong scene. For each correctly-answered question we identified the time points in the movie that corresponded to the scenes described by the experimenter (in the setup to the question) and by the patient (when giving a correct answer to the question). We then identified the neural state patterns that were active during those movie-viewing time points and correlated them with neural patterns from the corresponding parts of the interview. For descriptions made by the interviewer, we added an additional 4 seconds of padding in the neural data after the end of the description; this was done to account for delays in processing and understanding on the side of the patient. For the patient’s answers, we considered neural data from up to 10 seconds of the patient’s speech for each answer (this decision was made because answers exceeding 10 seconds typically digressed to irrelevant information that no longer formed part of the correct scene description; importantly, the below-described findings of neural reinstatement are highly invariant to these parameter choices and other analysis decisions – for instance, whether to use Pearson or Spearman correlation). To test for the significance of pattern-reinstatement, we derived a null distribution from the correlation with different scenes in the movie to obtain a z-score and p-value for each channel (see Methods). Controlling the false discovery rate at *q*= 0.05, we observed significant evidence for reinstatement of the corresponding state-patterns during the interview on 53 channels *p*s *<*= 0.0017 (Figure 2b, smallest z-score: *z* = 2.930, significant channels per patient: mean = 5.3, S.D. = 7.513, range = 0–25, median = 3). This confirms that neural activity during the interview reflected reinstated patterns from relevant parts of the movie, an effect that cannot be explained by nonspecific properties of the states. For further analysis, we fit a Gaussian Mixture Model with 2 normal distributions to the z-scores to separate the data into channels with and without reinstatement effects (this is a more conservative alternative to thresholding based on the assumption of a standard normal distribution; see Methods). The distribution with the lower mean serves as an empirical estimate of the null distribution^31^. We then selected channels if their z-score was at least two standard deviations above the mean of the empirically derived null distribution. With those parameters, we obtained a threshold of *z* = 2.3274 (i.e., a stricter threshold than *z* = 1.96), resulting in 84 channels that were selected for further analysis (Figure 3a, Supplementary Figure 4), henceforth referred to as cortical reinstatement channels (CR-channels). These channels were located in brain regions that overlap substantially with the default mode network (DMN)^38^; 42 of CR-channels were located closest to the DMN in the 7 Network solution defined by Yeo and colleagues^39^, 16 further CR-channels were closest to the somatomotor network, 11 to the frontoparietal network, 6 to the dorsal attention network, 4 to the visual network, 3 to the ventral attention network, and 2 to were closest to the limbic network. The number of CR-channels falling in the DMN (42/84) was significantly greater than expected based on the ratio of electrodes located in the DMN (393*/*1406; exact binomial test, *p <* 0.001).

**Figure 3:**
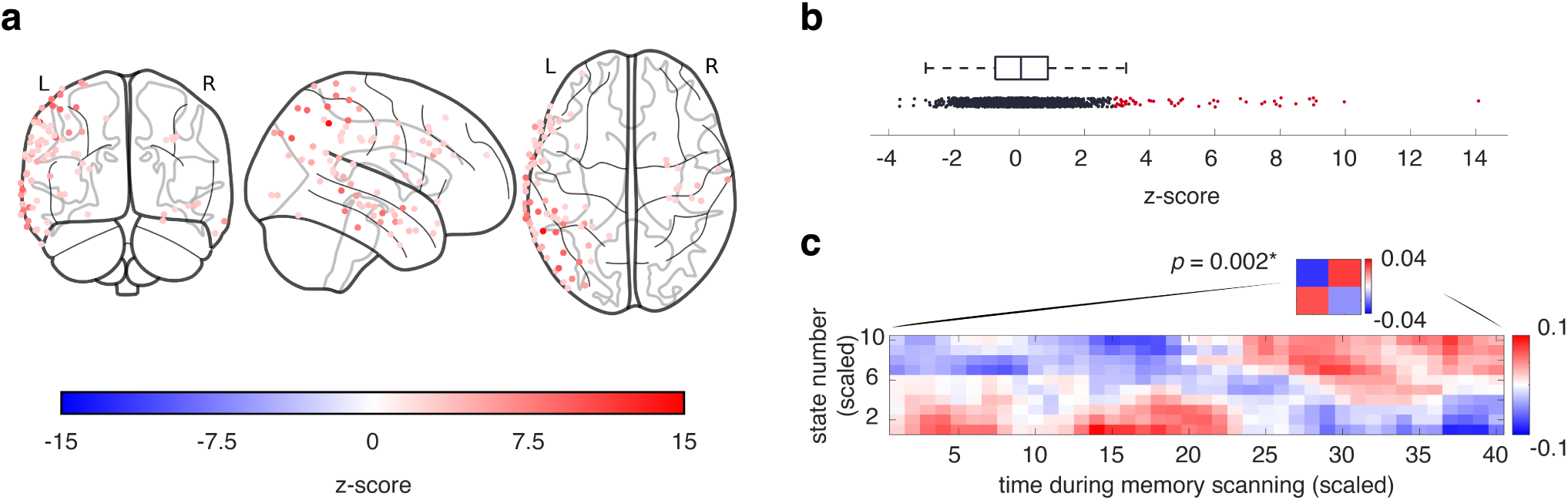
Neural states are reinstated during the naturalistic interview and unfurl in a forward direction during silent memory-scanning. **(a)** Cortical reinstatement channels (CR-channels), where evidence for reinstatement of state-patterns (during periods when the interviewer or patient were talking) exceeded the mean of the null distribution by 2 standard deviations. State-pattern reinstatement was observed in regions that overlap substantially with the default mode network (DMN); specifically, 42 of CR-channels were located closest to the DMN defined in the 7 Network solution defined by Yeo et al.^39^ (16 were closest to the somatomotor, 11 to the frontoparietal, 6 to the dorsal attention, 4 to the visual network 3 to the ventral attention, and 2 to the limbic network). **(b)** Z-score of evidence for reinstatement on all channels. Boxplot represents 25*^th^* and 75*^th^* percentile around the median, whiskers represent minimum and maximum excluding outliers. Black dots are z-scores on individual channels, red dots are z-scores where the corresponding p-value exceeds the threshold of significance from controlling the false discovery rate at *q* = 0.05. **(c)** Reinstatement of neural states during silent memory-scanning: The figure shows the average state-pattern by time-during-silent-memory-scanning correlation matrix, after it was scaled to 10 state-patterns and 40 time-points. The inset represents the same correlation matrix scaled to 2 state-patterns and 2 time-points. Statistical significance and direction of memory-scanning was assessed based on the 2 by 2 scaling. The correlation matrix illustrates that early state-patterns from the scanned segment correlate highly with early memory-scanning times while later state-patterns correlate highly with late memory-scanning times.

### Neural event states are unfurled in a forward direction during memory-search

We now tested if there was also significant evidence for reinstatement of state-patterns on CR-channels *during the memory search period* (when neither patient nor experimenter were speaking). This period starts after the end of the description of the setup and ends at the beginning of the answer period. For this analysis, we looked for reinstatement of all state-patterns that occurred (during viewing) in the period of time between the setup scene and the target scene (inclusive of the target scene, excluding the setup scene). Note that successive state-patterns from the movie-viewing phase can be uncorrelated or even negatively correlated with each other. Consequently, the average correlation with all state-patterns during a particular bout of silent memory scanning may average to zero or even to a negative correlation as multiple states are traversed. Nonetheless, when subjecting the average correlation with scanned state-patterns from the movie to a permutation test (randomly flipping the sign of correlations for each patient), we observed significant evidence for reinstatement during the scanning period (*p* = 0.0136). Next, we analyzed the order in which neural state-patterns unfurled by subjecting the average correlation between the first and second half of scanned state-patterns and the first and second half of memory-scanning to an ANOVA across 81 memory-scanning trials (Supplementary Table 2, compare also: Figure 3c). We found a significant main effect indicating unequal correlations (p *<*0.0022).

We then averaged Fisher-Z transformed correlations to compare the matching times (first half and first half, second half and second half) to the average of non-matching times (first half and second half, second half and first half); effectively testing forward vs. backward scanning. A positive difference supported by a significant post-hoc t-test (*t*(80) = 2.268, *p* = 0.013) confirmed that memory-scanning proceeded in a forward direction (compare Figure 3c, inset).

### Correlates of state transitions during memory-search in cortex

A central goal of our analyses was to look for fast neural processes occurring at moments of state transitions during memory-scanning. To identify those moments of state transition on CR-channels during memory scanning, we used Dynamic Time Warping (DTW)^40^ to compute the optimal alignment between neural state-patterns on CR-channels during movie viewing and the neural time series during memory scanning (Figure 4). In brief, for each trial of memory-scanning and each CR-channel, we computed the state-by-time correlation matrix and found the path from state 1 and time-point 1 (bottom left of the matrix) to the last state and the last time-point (top right of the matrix) that maximizes the overall correlation between state-patterns and data-points in the memory-scanning interval (note that DTW only allows for forward transitions on the warp-path). This path reflects the best state-pattern to memory-scanning alignment; time-points of state-transition on this path consequently identify the most likely moments of state-transition during memory-scanning (compare Figure 4b).

**Figure 4:**
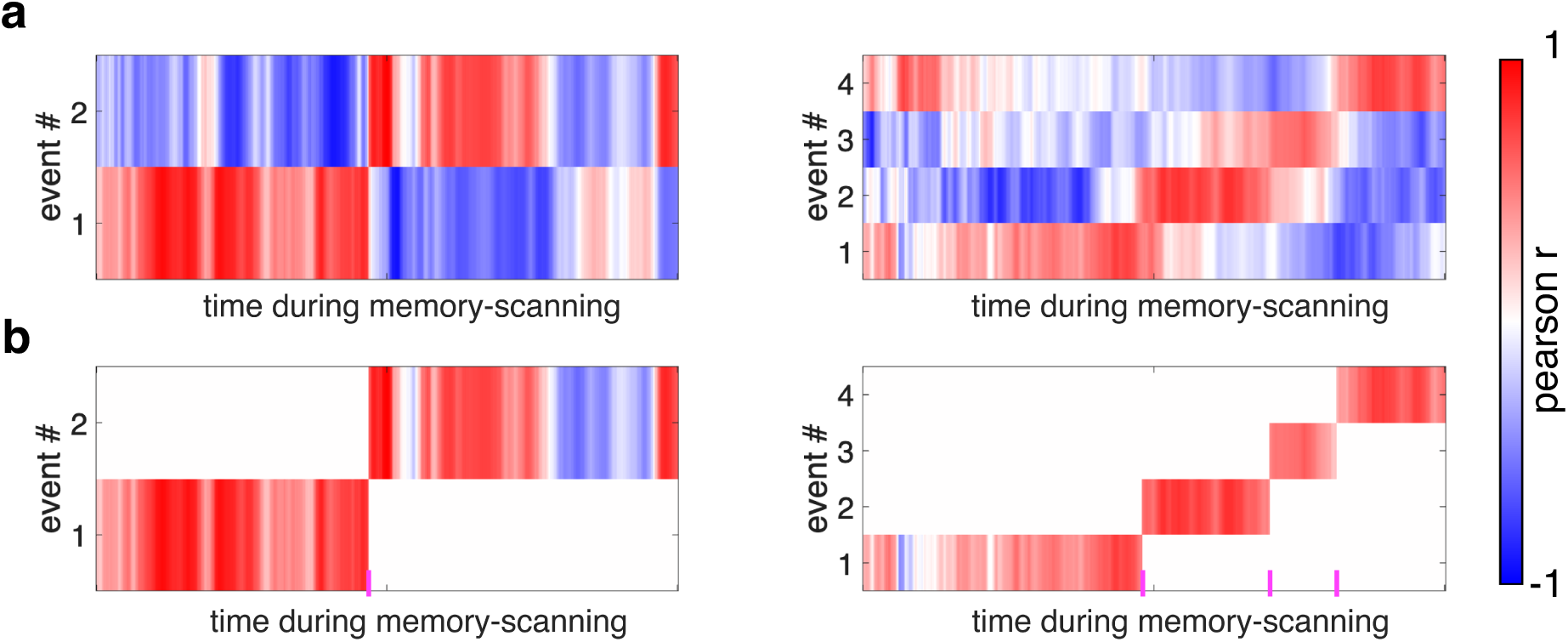
Detection of state transitions during silent memory scanning. **(a)** Event by time correlation matrices from the critical silent memory-search period where patients scanned their memory for an answer (see Figure 1). Left: example with one state transition in a trial where the patient searched for the answer to how Lieutenant Kowalski (George Clooney) wants to get back to earth. The memory-search period ends when the patient starts answering. Right: example with three state transitions as the patient searched for the answer to: “When do we see the space debris coming back?” **(b)** Optimal alignment path for data from (a) based on Dynamic Time Warping. Event transitions are indicated by transitions on the y-axis (marked on the x-axis with magenta lines). These are the points of event transition during silent memory-search that are investigated in subsequent analyses.

Based on these newly identified time-points, we next tested for univariate correlates of state-transitions during memory-scanning on CR-channels. As the first step, we computed the baseline-corrected Power Spectral Density (PSD, see Methods) for frequencies between 1 and 30 Hz and tested for a change in PSD at the exact time of state-transition. A two-tailed t-test comparing the relative PSD against zero revealed a significant decrease in the theta frequency band (4-6 Hz, *p*s *<*0.0002, controlling the false discovery rate at *q* = 0.025). We further tested for a cluster of changes in PSD within a window between 1 second prior and 1 second after the state-transitions during memory-scanning and observed a cluster of significant power decreases (Figure 5a) spanning a time-interval between 220 ms prior to the state-transition and 180 ms post state-transition and the frequency bands of 3-30 Hz (*p* = 0.035; two-sided).

**Figure 5:**
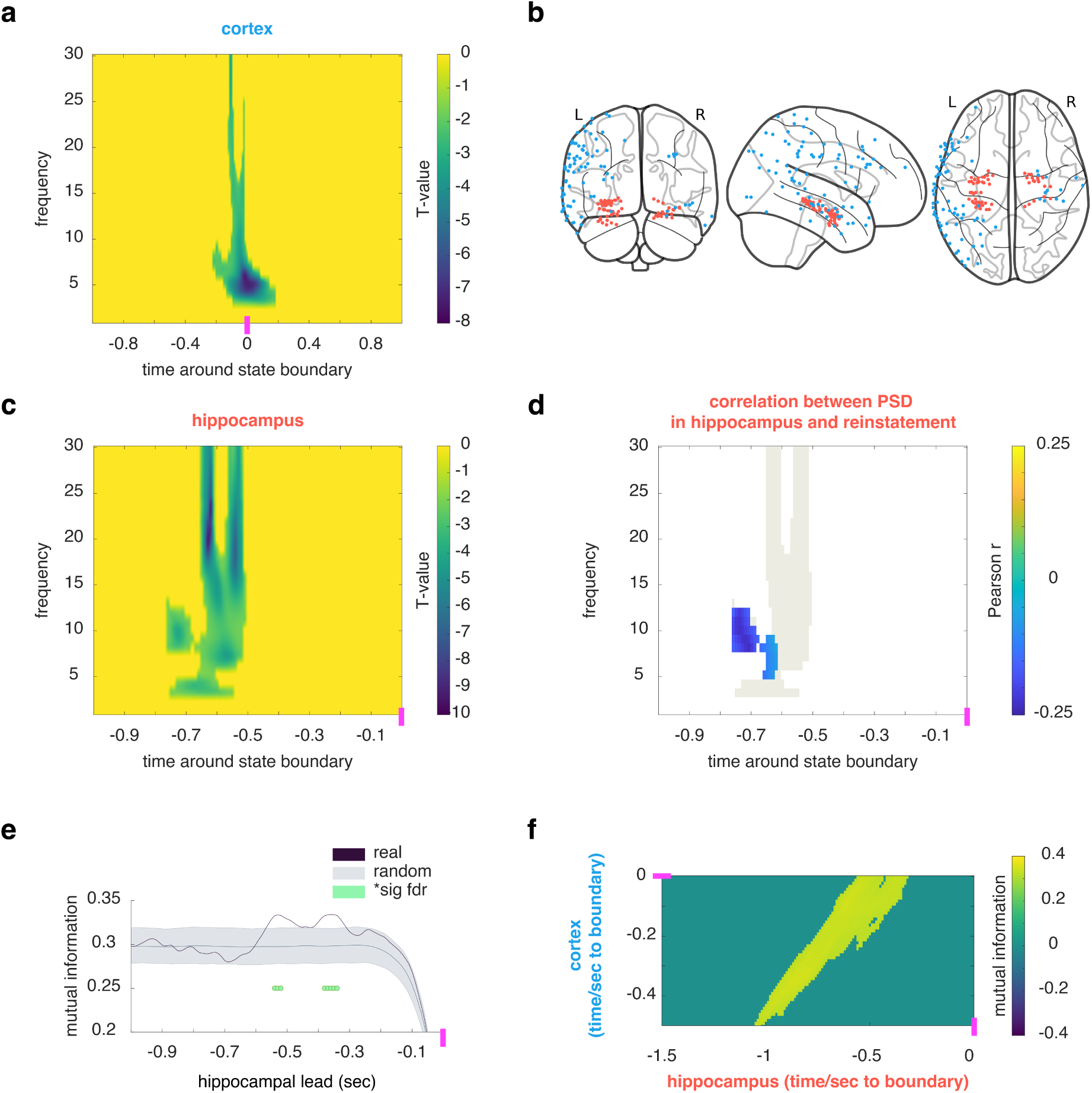
Neural underpinnings of state-transitions during memory-scanning. **(a)** Changes in Power Spectral Density (PSD) at state-transitions (magenta). A significant decrease in PSD was observed on cortical reinstatement (CR) channels in a cluster spanning 3-30Hz between 220 ms before and 180 ms after state-transitions. **(b)** CR channels (blue) and hippocampal channels (red). **(c)** A significant decrease in PSD was observed on hippocampal channels in a cluster spanning 3-30Hz between 760 and 510 ms before state-transitions (magenta). **(d)** Correlation between PSD in hippocampus (c) and strength of neural reinstatement of the upcoming state in CR channels (compare Figure 3c and Figure 4b). A significant negative correlation was observed in a cluster spanning 5-12 Hz between 760 and 619 ms, i.e., lower power before state transitions in hippocampus predicted more evidence for reinstatement throughout the upcoming state. **(e)** Evidence of information-flow from hippocampus to cortex at state transitions: Gaussian Copula Mutual Information (GCMI) in frequency bands 3-7, 8-15, and 15-30Hz between hippocampal channels and CR-channels, for real data (black line) and phase shuffled data (gray line SEM). GCMI is shown at different hippocampal leads (negative lag), conditioned on zero-lag GCMI. Green dots mark points where GCMI of the real data significantly exceeds GCMI for phase-shuffled data, controlling the false discovery rate at *q* = 0.05; the magenta line marks the state transition point. **(f)** The same analysis as in (e) at different times before the state-transition (magenta); y = 0 corresponds to the results in (e). The plot shows the largest cluster where GCMI between hippocampal and CR-channels in the real data significantly exceeded GCMI for phase-shuffled data.

### Correlates of state transitions during memory-search in hippocampus

We hypothesized that hippocampal neural activity would precede state-transitions in cortex and support the unfurling of a continuous memory trace. Testing this hypothesis, we computed the average baseline corrected PSD across hippocampal channels locked to each CR-channel’s state-transitions. We then tested for a cluster of changes in PSD up to the moment of state-transitions (starting 1s prior). We found a cluster of significant power decrease in hippocampus that spanned the time period from 760–510 ms before the cortical state-transition (Figure 5c) and extended across the frequency bands of 3 to 30 Hz (*p* = 0.022; two-sided).

### Hppocampo-cortical interactions underpinning state transitions

To test whether hippocampal power decreases were indeed related to the cortical reinstatement of state-patterns throughout the memory-scanning period, we next correlated the time-frequency bins inside the significant cluster of hippocampal PSD with the average amount of reinstatement that we observed for the state that the patient transitioned to next. Specifically, across all state transitions on all CR-channels, we correlated PSD in hippocampal time-frequency bins with the strength of neural reinstatement for the next (post-transition) state; strength of reinstatement was operationalized as the Fisher-Z transformed average correlation between the pattern of activity present during the next memory-scanning state and the pattern of activity from the corresponding state during movie-viewing (as identified by the dynamic time warping algorithm). By randomly flipping the sign of correlation within each patient, we identified a cluster of significant correlations (*p* = 0.02, one-tailed) where lower PSD in hippocampus was associated with more evidence for reinstatement of the upcoming state (spanning 5-12 Hz and from to 760 – 619 ms before state transitions; Figure 5d). In a control analysis, we correlated PSD before event transitions with the evidence for reinstatement of the current state (i.e., the state that is about to be abandoned in the state-transition), and we did not observe any significant effects (no cluster emerged, *p >* 0.999). Comparing these correlations with the upcoming state and the current state statistically, we further observed a significant cluster where the correlation with the upcoming state was stronger than the correlation with the current state (*p* = 0.015); this cluster emerged almost entirely within the cluster of significant association between PSD and reinstatement of the upcoming state (97.56% of the former cluster overlapped with the latter cluster, compare Figure 5d). These analyses confirm that the hippocampal power decreases prior to event transitions in cortex are linked to the reinstatement of upcoming state patterns.

Finally, we tested if there was information flow between hippocampus and cortex at state transitions: Across trials, does the amplitude of the hippocampal signal (in frequency bands 3-7, 8-15, and 15-30 Hz) at state transitions predict the amplitude of the cortical signal in CR-channels in those same frequency bands? Specifically, we computed the conditional Gaussian Copula Mutual Information (GCMI)^41^ between each hippocampal and CR-channel (Figure 5b) at various hippocampal lags (offsets). We conditioned GCMI on the hippocampal signal at zero-lag to account for volume conduction and common noise^31^. When comparing to GCMI derived from phase shuffled data, we observed significantly enhanced connectivity between hippocampal channels and CR-channels at two offsets (−540 to −520 ms and −380 ms to −340 ms) peaking at a 530 ms hippocampal lead and a 360 ms hippocampal lead (*p*s *<*0.0033, controlling the false discovery rate at *q* = 0.05); i.e., 3-30 Hz amplitude in hippocampus prior to state-transitions was significantly related to CR-channels’ 3-30 Hz amplitude at time of state-transition (Figure 5e). Finally, we wanted to account for the possibility that information from hippocampus arrives in cortex slightly before state-transitions. Therefore, we repeated above analysis with a sliding window starting 500 ms prior to the CR-channels’ state boundaries. When comparing patients’ average GCMI between hippocampal and cortical channels to GCMI from phase-shuffled data (up to an offset of 1-second hippocampal lead), we observed a cluster of significantly enhanced GCMI throughout the 500ms period that leads up to the state-transition (*p* = 0.008). This cluster spanned hippocampal leads from at least 550 ms (max = 580 ms lead) to leads of 280 ms (min = 530; Figure 5f); in sum, we observed information-flow from hippocampus to cortex prior to state-transitions and at state-transitions in cortex. This analysis of information-flow affords additional evidence that hippocampal power decreases are functionally related to cortical power decreases at and before event transitions.

## Discussion

These findings elucidate the slow and fast neural mechanisms that support memory-search across extended periods. By using a time delay embedding transformation^35, 36^ of iEEG data, we were able to identify spatio-temporal patterns in the default mode network (DMN) of neocortex that persisted over long periods during movie viewing. Those patterns correlated with behavioral measures of event perception and were reinstated throughout a naturalistic interview, where they unfurled in a forward sequence during memory-scanning. Computing the optimal fit between neural states during viewing and memory-scanning allowed us to pinpoint the exact moments when one event ended and the next one began during the memory-scanning period. Time-locking our analysis to these neural event boundaries then allowed us to test our predictions about the fast neural mechanisms that support memory search. Indeed, we found that low-frequency power decreases in the DMN marked the exact moment of state-transitions. Moreover, state-transitions were preceded by power decreases on hippocampal channels by approximately 700ms. Hippocampal power decreases further predicted the strength of memory reinstatement for the upcoming state, but not the previous state, and analyses of mutual information between the power-spectra in the hippocampus and cortex suggested that information was transferred from hippocampal to cortical channels at and prior to the state transitions.

Many features of these results align with and build upon existing findings in the literature relating to slow and fast memory-related processes. Our finding that behavioral annotations of event boundaries aligned with transitions in slow states during movie-viewing (measured using iEEG) is a conceptual replication of prior work that used fMRI^19, 21^, but adds a new level of temporal precision. Likewise, our finding that reinstatement of these slow states occurred in the DMN during the interview (when the interviewer or the patient were talking) fits with a large number of fMRI studies that have also observed reinstatement of event representations in the DMN^19, 22, 23, 25, 42–46^. With regard to fast memory-related processes: Our finding of decreased power spectral density (PSD) at state transitions during memory-scanning is consistent with numerous other electrophysiology studies (using other paradigms) that found an association between decreased PSD and memory reinstatement^13, 47–50^.

Apart from these similarities, there were also important differences between our study and existing work. Prior neural studies that explored memory processes at event boundaries used paradigms where the event boundary could be clearly marked based on stimulus properties or participants’ behavior^19, 21, 23, 25, 31, 44, 44, 51–55^. Our study, by contrast, sought to characterize hippocampus-cortex interactions during periods of silent memory scanning – a fundamental memory process that previous neural studies had not examined. The key innovation that made this possible was to use slow-timescale analyses (identifying moments of state transition) to time-lock the fast-timescale analyses of the neural correlates of transitions from one state to the next. In the absence of the state-transition information gleaned from the slow-timescale analyses, there would have been no way to determine when during the silent memory-search period to look for neural correlates of these transitions, which occur at a millisecond timescale. Time-locking to state transitions revealed a rich array of hippocampal and cortical dynamics occurring during this previously-unstudied silent memory search process. In addition to PSD decreases (in both hippocampus and cortex, with hippocampus preceding cortex), we crucially found relationships between hippocampal and cortical activity, whereby hippocampal PSD decreases predicted cortical reinstatement of the upcoming event, and there was time-lagged mutual information between early hippocampal and later cortical activity. These relationships provide support for our hypothesis that hippocampus supplies cortex with retrieved information at state transitions during silent memory search, bridging between adjacent events. Note that, in a memory-scanning task like ours, a failure to “bridge” from one event to the next can potentially cause the memory-scanning process to fail (whereas a failure to “bridge” during movie watching or story listening will, at worst, lead to a failure to anticipate upcoming events). A limitation of our study was that errors were too variable between patients to meaningfully analyze differences between correct and incorrect responses; future work can explore whether failures in hippocampal retrieval at state transitions during memory scanning ultimately lead to failures to locate the sought-after event.

In conclusion, the present study provides new insights into the slow event dynamics and fast hippocampal-cortical interactions that support memory search. In our prior behavioral work using this paradigm^14^, we inferred that event boundaries serve as “access points” for memory retrieval during continuous memory scanning based on careful analysis of reaction times (e.g., by showing that memory search times are better predicted by the number of events in the scanned period and, crucially, the distance of the search target to the previous event boundary than by the clock-time durations of those events; see also^15–18, 56, 57^). Here, through the direct time-resolved tracking of state-transitions in iEEG during continuous memory-scanning, and the concurrent recording of hippocampal activity, we were able to directly observe the neural mechanisms underlying memory-search. At these moments of state-transition, we were able to show transfer of information from hippocampus to cortex and a correlation between hippocampal activity and subsequent cortical reinstatement, thereby demonstrating how the hippocampus acts at state transitions to support the unfurling of temporally-extended memories.

## Data availability

Pre-processed data to reproduce the analysis in this manuscript is available in the Zenodo repository https://doi.org/10.5281/zenodo.15022125

## Code availability

All analysis code used in this manuscript is available in the Zenodo repository https://doi.org/10.5281/zenodo.15022125

## Acknowledgments

This work was supported by R01 MH112357 awarded to U.H. and K.A.N., S.M. was funded by Deutsche Forschungsgemeinschaft project 437219953. The authors thank Chris Baldassano for helpful comments and discussions.

## Online content

### Methods

#### Participants

A sample of 203 participants was recruited in online experiments for the purpose of norming the stimulus material for event boundaries. This norming experiment was previously published as part of a behavioral study. From 203 collected participants, 33 data sets were excluded for the first movie and 28 data sets were excluded for the second movie because of lags in stimulus presentation. Further details on recruitment, compensation, and exclusion criteria can be found in^14^. A distinct sample of 10 intracranial electroencephalography (iEEG) patients (18 − 41 years old, *mean* = 29.3, *SD* = 13.236, 4 male, 6 female, 9 right-handed, 1 ambidextrous) was tested at the Comprehensive Epilepsy Center of the New York University School of Medicine. Patients had been diagnosed with medically refractory epilepsy and were undergoing intracranial recording for medical purposes. The Princeton University Institutional Review Board granted ethical approval for all studies, additionally, the Institutional Review Board at the New York University Langone Medical Center granted ethical approval for the iEEG studies.

#### Stimulus material

To generate the *video material*, the movie Gravity (2013, Alfonso Cuaron) was edited to tell a coherent story of 15 min duration. This edited version resembles an extended trailer spanning the whole movie and features many of the key scenes in the story; for the experiments, it was divided into two halves of 7 minutes and 30 seconds duration. Interested readers can view the original movie for reference or contact the authors to view the edited versions, however, we cannot publicly share the video material due to the copyright. A set of 18 *memory-scanning questions* was used to induce forward memory-scanning; those questions had been piloted and tested for difficulty in previous behavioral experiments^14^. The questions follow a specific format, they consist of a setup, and a target: The setup describes a scene (Scene A) in the video with the purpose of orienting the patient to that scene, e.g., “[In the space station] we see little flames flying […] into the hallway”; the target part of the question is typically introduced by “When is the next time…” and asks about a later scene in the video (Scene B), e.g., “[…] that we see fire?” (compare: Figure 1). This “when is the next time…” framing was meant to induce participants to scan forward in memory from the setup to find the *next* instance of the target (e.g., fire), instead of attempting to retrieve *any* instance of the target (of which there were often multiple instances; e.g., multiple times when something caught on fire).

#### Experimental procedures

The norming of the stimulus material is described in detail in^14^. In brief, online behavioral participants viewed the videos using their internet browser under the instruction to press the space bar whenever, in their opinion, one natural and meaningful unit ended, and another began. Thereby, each participant indicated at what times during the movie they perceived an event boundary^6, 7^. For the *naturalistic interviews*, iEEG patients were not asked about (and did not receive any information about) event boundaries. Instead, they performed an interview task that prompted them to perform memory-scanning. Upon giving informed consent to participate in the experiment, patients received comprehensive instructions from the experimenter. Here, the experimenter explained the tasks and emphasized the necessity for careful attention as patients watched the videos. The experimenter further asked the patients to provide vivid and detailed descriptions when they answered the memory-scanning questions. Prior to starting the experiment, patients also received additional information about the movie’s main characters and locations. These details were read aloud and ensured that patients could effectively follow the story line and were familiar with less common terms used in the movie (e.g., the Soyuz spacecraft and the Hubble telescope). Next, the patients engaged in a practice session with a brief clip of 9 seconds duration that featured Charlie Chaplin. This practice served to familiarize patients with the experimental procedure and with the format of the subsequent main movie-viewing and interview phase; if needed, the practice session was repeated.

Patients then started the main experiment by viewing the first half of the movie, which was then followed by the memory-scanning interview. During the interview, the first set of 10 memory-scanning questions that pertained to this video was asked in randomized order: First, the patient was told that they were about to hear a new question. The experimenter proceeded to read the setup of the question, which asked the patient to focus on a particular part of the movie (scene A); the experimenter then confirmed with the patient that they remembered that moment. If the patient confirmed, the experimenter proceeded to ask about a later part of the movie (scene B) by reading the target part of the question. After the patient described that scene in their answer, the experimenter proceeded to the next question. When all questions had been asked, the experiment continued with the second half of the movie, upon which another set of 8 interview questions were asked in randomized order. The reading of interview questions at the bedside in free interaction with the patient is a loosely structured task that requires a high amount of guidance and intervention from the experimenter: In some cases, the patient did not remember the setup part of the question; the experimenter could then try to describe the scene in more detail and give additional context to help the patient find the correct scene in memory. Once the patient did remember the correct scene (scene A) and was therefore oriented, the experimenter would proceed by asking about the target to elicit memory-scanning. If the patient could not remember the setup at all, the experimenter would proceed with the next question. Sometimes, the patient would give a brief answer that did not provide sufficient detail to unambiguously determine if they were referring to the correct scene (scene B); in that case, the experimenter could ask the patient to provide more details, which was then used to determine whether their previous answer was describing the correct scene. In the latter scenario the experimenter would often follow up with a reminder, asking the patient to provide rich detail in their future answers. Finally, the patient would sometimes describe more than the correct scene (scene B) in their answer and proceed describing subsequent parts of the movie in chronological order. In that situation the experimenter could interrupt the patient and thank them for providing the correct answer. Once the interview process was completed, patients were debriefed and thanked. They were given an opportunity to ask questions or voice any concerns. Due to experimenter error, two patients received the interview questions in the same order. For one patient, a question relevant to the second half of the movie was inadvertently posed after the first half but was then repeated in the second interview. In another instance, a question pertaining to the first segment was asked after the second; in that specific instance, the question was excluded from subsequent analysis.

#### Data collection

In the patient studies all video material was presented on a 15-inch MacBook screen using 85% of the screen space against a black background; throughout the entire experiment, high-quality audio recording was continuously maintained, capturing every verbal interaction in the room. The onset of the two movie stimuli was recorded with a video-trigger through a photo-diode. The diode was attached to the bottom-left corner of the screen on the patient laptop and connected to a channel of the clinical amplifier that concurrently recorded intracranial EEG recordings from the patient. Before the start of the movie, a white rectangle was flashed underneath the photodiode for a duration of one second and then set to black for an additional second three times. For the duration of the movie the rectangle was then set to white again, i.e., the fourth onset of the photo diode marked the onset of the movie. When the movie ended, the rectangle under the photo diode was turned off again and flashed three more times with a one second duration and interval, marking the end of the video presentation; the flashing of the trigger in known intervals served to validate the accuracy of timing for on- and offsets. Additionally, an audio cable was connected from the laptop to the iEEG amplifier and an audio pulse was sent concurrently with each onset of the video-trigger.

Neural data were collected via grid arrays (8 × 8 contacts, 10 or 5 mm spacing), linear strips (1 × 8/12 contacts), depth electrodes (1 × 8/12 contacts), and a high-density grid (16 × 8 contacts, 1 mm spacing) for one patient over the lateral posterior occipital cortex. The recording was realized with a NicoletOne C64 clinical amplifier (Natus Neurological, Middleton, WI), data were referenced online to a two-contact subdural strip close to the craniotomy location. Data underwent filtering through an analog bandpass (0.16–250 Hz range), they were digitized at 2048 Hz and down-sampled to 512 Hz. Available data spanned 1515 channels (ranging from 104 to 242 channels per patient, detailed in Supplementary Figure 1). Out of these, 109 channels were omitted from the analysis due to insufficient signal quality (varying between 1 to 33 channels for each patient, as detailed in Supplementary Table 1).

**Supplementary Table 1:** Characteristics of the patient sample and details on the number of analyzed channels.

| patient # | sex<br>(m/f/d) | hand | diagnosis | N channels<br>(raw) | N channels<br>(included) | HC channels<br>(included) | CR channels |
| --- | --- | --- | --- | --- | --- | --- | --- |
| 1 | m | ambi | treatment-resistant focal epilepsy arising from left hemisphere, not clearly localized | 161 | 141 | 7 | 28 |
| 2 | f | right | treatment-resistant focal epilepsy arising from left hemisphere, not clearly localized | 120 | 113 | 7 | 2 |
| 3 | f | right | drug-resistant focal epilepsy likely arising from left temporal lobe | 242 | 218 | 3 | 14 |
| 4 | m | right | drug-resistant focal epilepsy arising from left temporal lobe | 124 | 119 | 7 | 8 |
| 5 | f | right | drug-resistant non-lesional left temporal (likely neo-cortical) epilepsy | 133 | 131 | 3 | 4 |
| 6 | f | right | drug-resistant focal epilepsy, with focal aware and impaired aware seizures | 196 | 193 | 13 | 8 |
| 7 | m | right | focal epilepsy | 128 | 95 | 6 | 0 |
| 8 | f | right | refractory focal epilepsy and anxiety/depression | 132 | 131 | 4 | 2 |
| 9 | m | right | drug-resistant lesional focal epilepsy | 104 | 99 | 6 | 3 |
| 10 | f | right | drug-resistant focal epilepsy, likely arising from a right parietal lesion | 175 | 166 | 13 | 15 |

#### Audio data processing and alignment

To analyze the iEEG recording in conjunction with the interview data and the videos, several alignment and processing steps were applied. Using Audacity® (version 2.4.2, audacityteam.org), the high-definition audio from the patient room was first converted to mono and down-sampled to 16 kHz. Subsequently, the audio trace from the video stimuli was extracted at the same sampling frequency. Cross-correlation analyses between each video’s audio trace and the room recording were then computed in Matlab (2020b, mathworks.com). Specifically, the cross-correlograms between the audio time-series were computed and smoothed with a gaussian kernel (width: 32 sample points); the lag of the peak in the respective smoothed cross-correlogram was used as a marker of the exact onset of the videos in the audio recording. Next, the audio recording was used to manually transcribe the interview. Then, all words in the interview and during the movie were time-stamped via Penn Phonetics forced aligner^58^. To ensure precision, a custom-written graphical user interface was used to hand-select small sub-sections of text and their corresponding sub-sections in the audio ^1^, which effectively mitigated word alignment error due to ambient noise or extended pauses. The audio recording’s time scale was henceforth adopted as the reference time-scale for the experiment in all recordings, notably including the iEEG recording, where the onset of the movie was determined via the trigger pulses of the photodiode. In brief, the outcome of the procedure was the alignment between word onsets in the interview, the movie onset during the experiment, and the iEEG recording.

**Supplementary Figure 1:**
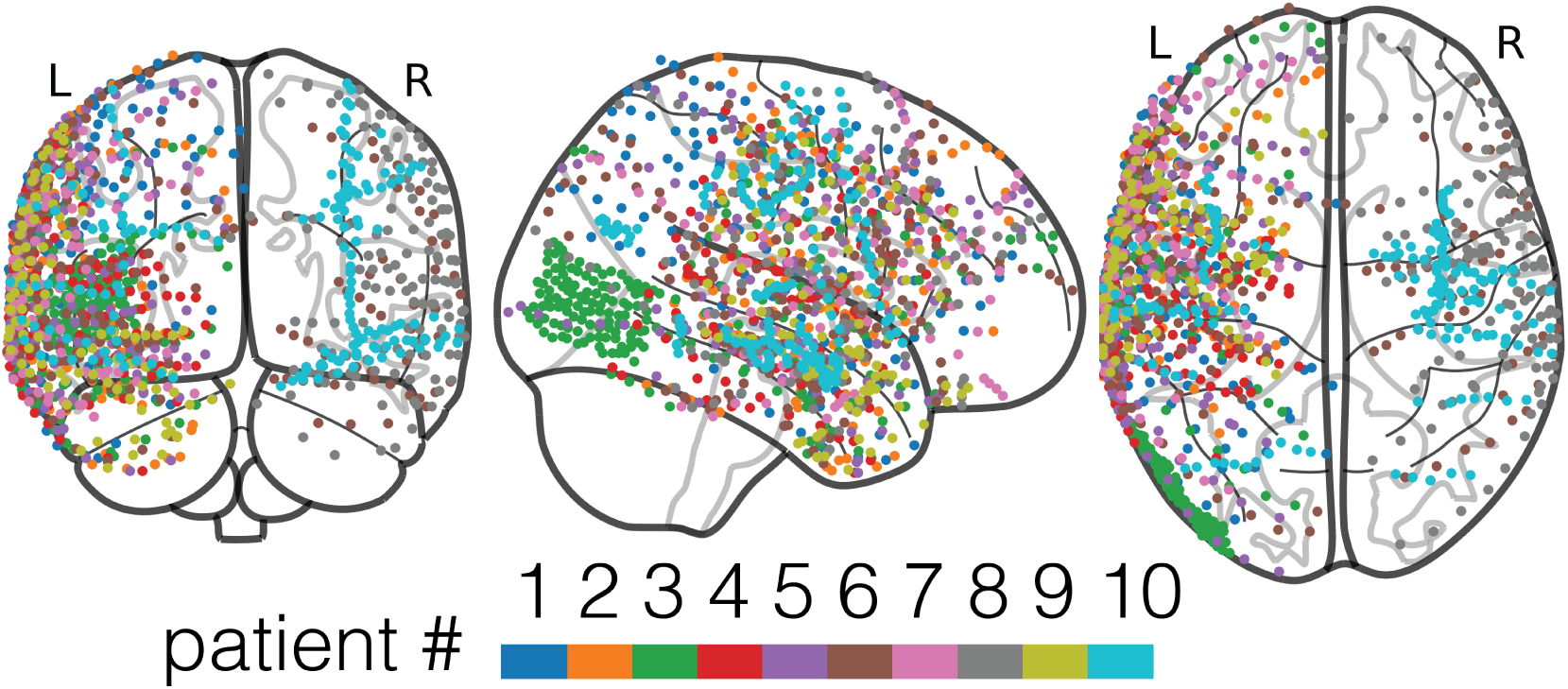
Position of analyzed electrodes. Electrodes from the same patient are coded in the same color.

#### Neural data processing

##### Electrode localization and alignment

Electrode localization was achieved by aligning either the post-surgical T1-weighted MRIs (patient 1 and 2), or post-surgical CT scans (patient 3–10) to the pre-surgical MRI of each patient^59^. Manual examination in MRIcroGL^60^ was performed to identify the hippocampal electrodes. Subsequently, nonlinear mappings of the MRIs to the Montreal Neurological Institute’s MNI-152 template were computed to transform electrode coordinates into MNI space.

##### Pre-processing

Electrophysiological data were pre-processed using the FieldTrip toolbox^61^ and custom written scripts in MATLAB (2020b, mathworks.com).

###### Alignment and segmentation

Triggers marking the start of each movie were used to synchronize iEEG recordings with the movie stimuli. To mitigate drift between recordings, the iEEG data were duplicated and each duplicate’s time-axis was aligned separately with the audio recording’s time-axis from the first and second movie, based on their respective triggers. The first duplicate was then cut to encompass the interval from four seconds before the first movie to five seconds before the second movie; the second duplicate spanned from four seconds before the second movie’s start until the end of the experiment. This process yielded two non-overlapping recordings corresponding to the first and second halves of the session.

###### Artifact Identification

Residual epileptic spikes and artifacts were marked with a semi-automatic procedure: Candidate artifacts exceeding 5 inter-quartile ranges above a channel’s median were marked along with 300ms around these values. Subsequently, only instances of artifacts where at least 3 neighboring channels concurrently identified them were retained, Finally, all potential artifacts underwent manual inspection and correction.

###### Re-referencing and interpolation

An ICA was computed to remove shared noise across channels^62^. ICA filters were generated using data from both experiment parts, previously filtered by a band-stop filter (stopbands: 55–65, 115–125, 175–185). The ICA computation was exclusively performed on artifact-free data. Next, raw data around manually revised artifact moments, where channel amplitude additionally exceeded 3.5 interquartile ranges above its median, underwent Monotone Piecewise Cubic Interpolation^63^ within a 150ms window. The previously computed ICA solution was then applied to the resulting data and maximally ten spatially broad components capturing reference noise were rejected (mean = 2.5, median = 2 components). Finally, a band-stop filter (stopbands: 58–62, 118–122, 178–182) removed residual line noise from the interpolated and re-referenced data. All filter operations were conducted using zero-phase lag 4th order Butterworth IIR filters.

##### Slow component analysis

###### Temporal embedding searchlight

Electrophysiological recordings exhibit intricate temporal dynamics, encompassing both fast and slow neural processes that manifest in complex spatio-temporal patterns of activity. To effectively leverage rich spatio-temporal patterns, we employed a time delay embedding (see Vidaurre et al.^36^): By concatenating lagged versions of the data along the channel dimension, we obtained spatio-temporal patterns throughout the recording^35^. To preserve spatial specificity, however, we limited this temporal embedding to 5 neighboring channels at a time, effectively combining the temporal-embedding approach with a searchlight procedure. Specifically, in each local neighborhood, we concatenated the data at 128 lags at a sampling rate of 512 Hz, thereby covering a window-width of 250 milliseconds (Supplementary Figure 2a).

###### Dimensionality reduction

We next performed a dimensionality reduction on these high-dimensional data. In contrast to Vidaurre et al^36^, who performed a principal component analysis (PCA) of the embedded data in their study, we used a novel dimensionality reduction procedure that was designed to emphasize slow components, i.e., components that maximize autocorrelation rather than variance (Supplementary Figure 2b). Our goal was to extract components that display the slow dynamics described by Baldassano et al (2017)^19^, staying stable for long periods of time within events and then shifting to a new pattern. To accomplish this goal, we computed the average data covariance between the embedded iEEG data and time-lagged versions of itself, at all lags between 252 milliseconds (1 sampling point outside of the temporal window of the feature vector, i.e., starting at lag k = 129) and 10 seconds (maximum lag, L = 5120 sampling points). Eigenvectors of this covariance matrix capture components of maximal covariance between the embedded signal and its shifted copies without favoring any specific frequency. We constrained these components to have a maximal covariance of 1, which is equivalent to computing a canonical correlation analysis (CCA) between the embedded signal and its own shifted versions (CCA maximizes covariance between two time-series X and Y while constraining the projected data to unit variance). Conceptually, this analysis performs a normalized PCA on the spatiotemporal patterns, where temporal jitter is introduced into the component estimation to ensure that components reflect stable patterns. This was done separately for the recording corresponding to the first and second video, i.e., no interview data was used in the computation of the slow component transformations.

In practice, we computed these components by solving the generalized eigenvalue problem:

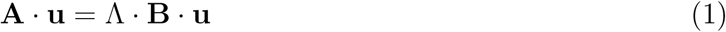

with 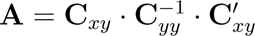 and **B** = **C***_xx_*.

**C***_xx_* and **C***_yy_* are covariance matrices of the signal without shifts; **C***_xx_* is computed with the signal truncated at the right end:

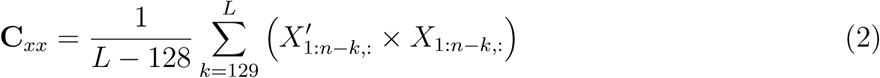

with k referring to the offset in the signal and L denoting the maximal number of lags (here: 10 seconds, i.e., 5120 points at a sampling rate of 512 Hz); n denotes the maximum number of sample points in the recording. **C***_yy_* is computed with the signal truncated at the left end:

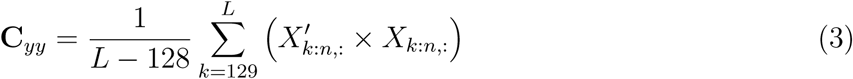

**C***_xy_* refers to the cross-covariance matrix between copies of the high-dimensional signal and its shifted versions.

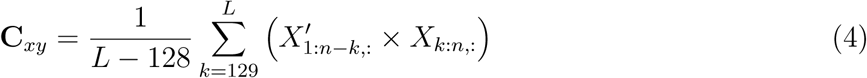

To increase stability, we additionally applied shrinkage regularization^64^ to all covariance matrices before computing the eigenvalue decomposition via:

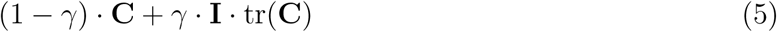

with a regularization parameter of *γ* = 0.0001.

To obtain a threshold for the retention of components from this analysis, we next com<u>p</u>uted the correlation coefficient of each component from the square root of the eigenvalues 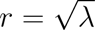 and derived a p-value as *p* = 1 − *tcdf* (*t, n* − 2), where tcdf denotes the cumulative distribution function of the t-distribution and t denotes the t-statistic 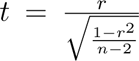. We retained components where the obtained p-value was smaller than 0.05; in cases where more than 300 components met this threshold, we only used the top 300 components.

For data *X_A_* and *X_B_* corresponding to the first and second video, we obtained eigenvectors *u_A_*and *u_B_*. Slow components could subsequently be computed by projecting the embedded signal via the computation of *X × u*. To avoid over-fitting, we projected each signal with the solution from the other movie (Supplementary Figure 2c); i.e., we projected *X̂_A_* = *X_A_* × *u_B_* and *X*^^^*_B_* = *X_B_* × *u_A_*.

###### Neural state segmentation

We performed a segmentation on the slow components that were extracted in each channel’s local neighborhood. First, we down-sampled the components to a sampling rate of 40 Hz for reasons of computational efficiency. We then applied greedy state boundary search^20^ to the data (with a block-size of 40 and a maximal number of 250 states). In 3 instances the algorithm did not converge on a solution (2 channels of subject 9 during the first video and 1 different channel during the second video); no further analyses were performed on these channels. From this analysis, we obtained neural states for each local neighborhood characterized by corresponding state-patterns. Furthermore, we obtained the boundaries between those states to a precision of *<*25 ms.

###### Representational similarity between movie viewing and parts of the interview where the experimenter or participant were speaking

To test for the reinstatement of neural state-patterns during the parts of the interview where the experimenter or the participant were speaking, *excluding* the silent memory-scanning period, we performed a representational similarity analysis (RSA) between state-patterns at encoding (during the presentation of the movie) and the slow component data during the interview. For this analysis, we correlated the components throughout the time period in which a given scene was described in the interview (either by the researcher or by the patient) with all neural state-patterns that corresponded to that scene in the movie during viewing. For the movie-viewing period, corresponding state-patterns were selected for correlation if the first movie-frame or the last movie-frame of the described scene was inside the interval spanned by the beginning and end of a neural state. Time intervals during the interview were selected for correlation based on the onset time of the first word in a description and the offset time of the last word in a description. To account for a delay in processing (the patient may take some time to understand which scene is being described), we considered an additional 4 seconds of data after the interviewer’s description (setup) in the interview. For answers, we considered the first 10 seconds after the onset of the patient’s response (later parts of the response often digressed into irrelevant content and were characterized by long pauses). We then computed the average correlation (averaged over time) between neural patterns from the interview and each state-pattern from encoding. We then compared the average correlation of corresponding intervals (same scene in perception and memory) to average correlations of non-corresponding intervals (different scene in perception and memory). Specifically, we averaged all correlations of same content (*r_same_*) and all correlations of different content (*r_diff_*). We further bootstrapped the standard deviation of averages (*SD_avg_*) by resampling 100,000 times from all computed correlations (corresponding and non-corresponding); crucially, these samples were of the same size as the number of corresponding correlations (*N_same_*). From this we computed a z-score: *z_rsa_* = (*r_same_ r_diff_*)*/SD_avg_*. Estimating the standard deviation of averages with the same number of correlations (*N_same_*) is important because averages across a larger number would underestimate the true standard deviation^47^.

**Supplementary Figure 2:**
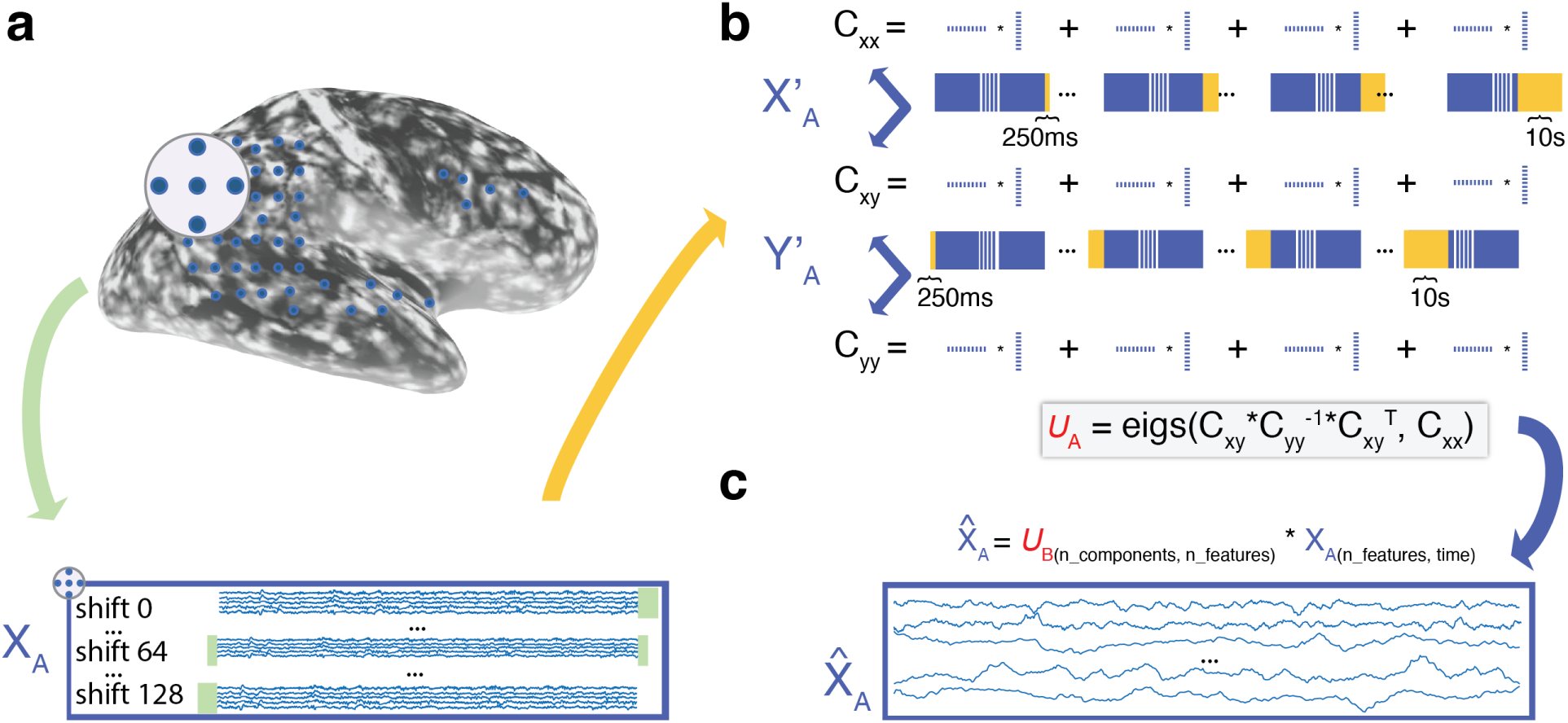
Slow component analysis. **(a)** A searchlight of 5 neighboring channels was used. Time courses were concatenated along the channel dimension for shifts up to 128 sample points (250 ms), effectively bringing the spatial and the temporal dimension together in spatio-temporal feature vectors. **(b)** Spatio-temporal feature vectors were mapped onto themselves concurrently at various lags. Specifically, the transformation matrix *U* maximizes the correlation of the signal with itself at lags between 250 ms and 10 s. The transformation was applied in a cross-validated fashion: The transformation fit to the first half of the data was applied to the data from the second half and vice-versa. **(c)** Projected feature vectors from (a) capturing spatio-temporal patterns that are stable over long periods of time.

###### Channel selection for further analysis

A substantial number of channels survived the correction for multiple comparisons, statistically confirming the existence of reinstatement. Crucially, however, many channels that did not survive the stringent correction for multiple comparisons nonetheless displayed high z-scores, suggesting that they might contain useful signal relating to reinstatement. For further analysis, we selected those channels on which the evidence for reinstatement exceeded at least two standard deviations above the null distribution. We derived that null-distribution empirically by fitting a Gaussian mixture model with two Gaussian distributions onto the z-scores from all channels; intuitively, the distribution with the lower mean is meant to capture noise and the distribution with the higher mean is meant to capture signal. All channels whose z-score was higher than 2 standard deviations above the mean of the lower distribution were included in subsequent analyses; this resulted in an empirical cutoff of z = 2.326, i.e., a value that is slightly more conservative than we would have obtained from assuming a standard normal distribution.

###### Representational similarity between movie viewing and silent memory-scanning during the interview

To assess reinstatement of event patterns in the *memory-scanning period*, we selected corresponding neural state-patterns from the scanned segment during movie-presentation. Specifically, state-patterns from the viewing period were selected starting with the first movie-frame after the scene described in the setup, and ending with the last movie-frame from the target scene. Memory scanning intervals were determined based on the offset time of the last word in a description of the setup and the onset time of the first word in the patient’s answer. We obtained z-scores again by comparing corresponding correlations (i.e., correlations to the state-patterns from the scanned segment of the movie) and non-corresponding correlations (correlations to other state-patterns, from outside of the scanned segment). We then computed a standard deviation by bootstrapping averages from all combinations of corresponding and non-corresponding correlations to derive z-scores.

###### Assessing the direction of memory-scanning

To assess the direction of memory-scanning, we analyzed the correlation of each neural state-pattern from encoding with each moment during memory-scanning, i.e., we derived time-courses of correlation for each neural state-pattern. Because each memory-scanning trial could span a different number of states, and each channel could have a different number of neural state-patterns, we aggregated the correlation time-courses in the following way: First, we discarded trials at a given channel if that channel did not have at least two state-patterns. Second, we scaled the states’ correlation time courses to the same number of states and to the same scanning duration for all subjects, channels, and trials. Finally, we averaged the time courses of correlation across channels, obtaining an averaged time course for each interpolated neural state for each trial. This allowed us to obtain an averaged state-by-time correlation matrix that captures the similarity of early and later (interpolated) neural states to different moments during memory-scanning via their correlation.

###### Identification of neural state-transitions during memory-scanning

We used the algorithm behind dynamic time warping (DTW) to compute the optimal alignment between the state-patterns from encoding and the component time course during memory-scanning^40^; we computed this separately for each subject, trial, and channel (Figure 4). Concretely, we used the correlation of the multivariate patterns as a distance metric between every state-pattern and every time-point during retrieval. We then computed the optimal warp-path that maximizes the correlation in the transition through state-patterns from the beginning of the memory-scanning period to its end. Each transition in that optimal warp-path from one neural state-pattern to the next (i.e., a step on the y-axis in the cumulated pairwise correlation matrix) was taken as a neural state boundary during memory-scanning.

###### Time-resolved frequency spectra at neural state-transitions during memory-scanning

During memory-scanning, we computed the power spectral density (PSD) around neural state-transitions separately for each channel on which we observed reinstatement (see above for channel selection). Specifically, for a given channel, we epoched the data centered on the moment of neural state-transition and convolved the data with Morlet wavelets. To maintain maximal temporal resolution, we used 3 cycles of a frequency and computed the spectral power for 1 Hz frequency bins between 1 and 30 Hz in steps of 10 ms. We centered this analysis on the time points of neural state-transitions and computed the average power, which we baseline corrected by z-scoring on the baseline between −2500ms and 2500ms around the neural state-transitions. In another analysis, we epoched all available hippocampal channels in the same way and centered them relative to each available cortical channel’s state-transitions; after computing the PSD and performing baseline correction, we averaged across all hippocampal channels. This resulted in a hippocampal homolog for each cortical channel, analyzed with respect to that channel’s state-transitions.

###### Correlation between hippocampal PSD and cortical reinstatement

To assess the relationship between PSD in hippocampus and reinstatement during memory-scanning, we computed the baseline-corrected PSD in the same way as described above, were each cortical channel has a hippocampal homolog. On cortical channels, we then averaged correlations along the warp-path determined by dynamic time warping (see above). Specifically, for each state in which patients lingered for at least one sample-point during memory-scanning, we averaged correlations between the corresponding state-pattern and all sample-points that were assigned to that state on the warp-path for each CR-channel. We then computed the Fisher-Z transformation (i.e., atanh) of that average correlation as a measure of reinstatement of the upcoming state. Thereby, we obtained a single correlation value for each state-transition that we matched to a value of baseline-corrected PSD for each time-frequency bin inside the cluster of hippocampal power decreases. Note that this analysis was limited to those channels where at least one state-transition was present and to those states that were visited for at least one sample-point. We then computed the correlation between PSD and reinstatement across all state transitions and CR-channels for each patient.

###### Mutual information connectivity analysis

Connectivity analysis between cortical and hippocampal channels was performed by assessing multivariate Gaussian Copula Mutual Information (GCMI^41^) between the amplitudes of the filtered signal (compare^31^). GCMI is a rank-based, robust method that makes no assumptions about the marginal distributions of each variable and is insensitive to outliers; its estimate is a lower bound of the true mutual information^41^. Specifically, we filtered the data in 4 frequency bands with a zero-phase lag 4th order Butterworth IIR filter as implemented in the fieldtrip toolbox^61^: a lowpass at 3Hz, and bandpass filters between 3 -7Hz, 8-15Hz, and 15-30 Hz. We then computed the absolute of the Hilbert transform and down-sampled each channel’s continuous time-courses to 100Hz. For every channel on which we observed reinstatement (see above for channel selection), we then centered a 1-second wide window at the time points of neural state-transition and computed GCMI with every hippocampal channel. Therein, the computation of GCMI was repeated at all possible lags (the offset between hippocampus and cortex), starting with a 1-second hippocampal lead and ending at a 1-second hippocampal lag. Specifically, the conditional GCMI was computed to account for spurious (zero-lag) connectivity, i.e., the GCMI between a cortical and HC channel at lag L was conditioned on GCMI between them at lag 0. We then averaged the GCMI across all hippocampo-cortical channel-pairs, which resulted in a GCMI value for each lead/lag in every subject. Next, we repeated this analysis at different offsets to the neural state-boundary, i.e, we shifted the centering from 500ms before the neural state-transition to the moment of state-transition.

### Statistical testing

#### Association between neural state boundaries and behavioral event perception

To test if the neural event boundaries that were derived in a data-driven way captured behavioral button presses that we recorded in online experiments, we computed a behavioral response profile around neural event boundaries for each channel. To this end, we locked the time course of behavioral agreement (derived from the proportion of button presses in the behavioral sample) to the neural event boundaries at each channel; this gave us the average behavioral response to a neural event boundary at each channel. We then computed a null distribution of that response by randomly permuting the order of neural states 1000 times (keeping their length intact) at each channel and recomputing the average response. Subsequently, we z-scored the observed behavioral response profile to each channel’s neural boundaries based on the permuted data, i.e., we subtracted the average response to shuffled data and divided by the standard deviation across permutations. Note that under the null hypothesis, which states that there is no systematic association between neural event boundaries and behavioral response profiles, the z-scored response profiles should not be statistically different from zero. We next tested whether there was an overall statistical association between neural state boundaries and behavioral responses: We computed the average response profile for each patient (averaging the time-locked z-scored response profile across all channels) and subjected the average response profiles to a two-sided t-test across patients at every time point. Finally, we corrected the resulting t-values for multiple comparisons across time on the interval starting 2 seconds before neural state-transitions and ending 10 seconds after neural state-transitions by controlling the false discovery rate at a threshold of q = 0.05. We additionally confirmed the significance of the effect by computing a cluster permutation: We thresholded the t-statistic across patients at a t-value corresponding to an alpha threshold of 0.05 and computed the cluster sum of the largest contiguous cluster. We then flipped the sign of values randomly for each patient and re-computed the cluster 1000 times. Finally, we obtained a p-value as the ratio of random permutations where the empirically observed cluster sum was exceeded.

#### Neural state-pattern reinstatement

To assess the significance of pattern reinstatement during the parts of the interview when the interviewer or the patient were talking, we subjected the p-value corresponding to each z-score in the cumulative normal probability density function to a false discovery rate correction across all channels. Next, we selected a subset of channels for further analysis (see “Channel selection for further analysis” section), and we tested whether the average reinstatement *during silent memory-scanning* exceeded chance level on those channels. To this end, we randomly flipped the sign of all correlations at the subject level and recomputed the averages 1000 times (i.e., for each subject all correlations were either multiplied by 1 or −1 before correlations were averaged). From this permutation we obtained a p-value as the number of random averages that exceeded the true average.

#### Direction of memory-scanning

To test for the direction of memory-scanning, we scaled the time-courses of reinstatement to a length of 2 temporal units and scaled the number of neural states to a length of 2. We then performed a Fisher Z transformation of the values and computed a one-way ANOVA across all trials effectively testing if the four bins of state-to-scanning-time correlation were significantly different from each other. We then averaged the correlations that corresponded to scanning in a forward direction (1,1) and (2,2) and contrasted them with correlations that corresponded to scanning in a backward direction (2,1) and (1,1) via a 2-sided one-sample t-test.

**Supplementary Table 2:**
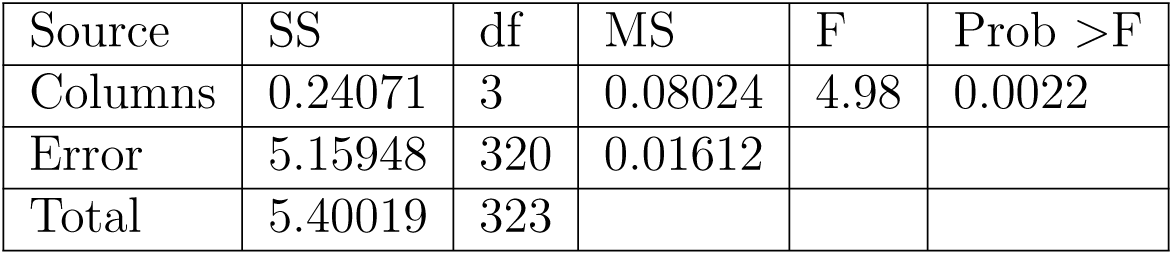
2x2 ANOVA across 81 memory-scanning trials.

| Source | SS | df | MS | F | Prob >F |
| --- | --- | --- | --- | --- | --- |
| Columns | 0.24071 | 3 | 0.08024 | 4.98 | 0.0022 |
| Error | 5.15948 | 320 | 0.01612 |  |  |
| Total | 5.40019 | 323 |  |  |  |

#### PSD and correlation between PSD and reinstatement

Changes in PSD at the exact moment of state-transitions in cortex were first tested against zero with a two-tailed dependent-sample t-test across 9 patients (for which memory-reinstatement was observed). This was done for each frequency bin. Subsequently, p-values were subjected to a correction for multiple comparisons across frequency bands by controlling the false discovery rate at *q* = 0.025. Changes in PSD at cortical and hippocampal channels were next tested with a cluster-based permutation test. Baseline-corrected power was first tested across 9 patients with a two-tailed t-test that tested PSD against zero in each time-frequency bin near state-transitions on CR-channels. For CR-channels, the statistical analysis was restricted to a time-window from 1 second prior to the state transition to 1 second after the event transition. For hippocampal channels (where we assessed whether neural correlates preceded state transitions), statistical analyses were restricted to a period from 1 second prior to the state transition to the moment of the state transition on the corresponding CR-channel. Neighboring t-values (falling into adjacent time or frequency bins) that exceeded the critical t-value at an alpha-level of 0.05 were subsequently summed across time and frequency bins, and the absolute of the sum within each cluster was computed. To create a null distribution, we repeated the same analysis 1000 times, but here we randomly multiplied each patient’s average PSD with 1 or −1. Finally, we compared the resulting null distribution of maximal absolute cluster-sum to the maximum sum that was observed in the real data and obtained a p-value as the proportion of instances that were more extreme in the null distribution. For the correlation values between PSD and reinstatement, we restricted the statistical analysis to the time and frequency period where a cluster of significant power decreases had been observed. Here we computed a one-sided dependent-sample t-test across 9 patients where we compared the Fisher-Z-transformed correlations against zero (as per our hypothesis of an association between power decreases and reinstatement, which yields a negative correlation). Again we compared the maximum cluster sum to the null distribution of cluster-sums under random sign-flip permutation across patients and report a p-value from the proportion of observations that are more extreme under random sign-flip.

#### Lagged mutual information

To obtain a null distribution of conditional GCMI, we generated surrogate cortical and hippocampal data by phase-shuffling their time-courses 1000 times using the method introduced in^65^. We then recomputed the exact same analysis for each of the surrogates. To test whether there was significant information-flow from hippocampus to cortex, we compared the GCMI at each lag between a 1 second hippocampal lead and a zero-lag, to the null distribution of GCMI that was derived from the surrogate data. Additionally, to obtain a higher numerical precision of p-values, we upsampled the surrogates by computing 100,000 averages of GCMI across patients, where we randomly selected one out of 1000 surrogate versions for each patient to compute the average^66^. We then controlled the false-discovery rate across lags to account for multiple comparisons. To test for significance on 2-dimensional maps of GCMI, we thresholded the average maps (averaged across patients for real and surrogate data) at the 95th percentile of the surrogate data. We then computed the sum of GCMI within each cluster and compared the distribution of maximum cluster sums in the surrogate data to the maximum cluster-sum in the actual data. We derived a p-value as the proportion of random permutations that resulted in a higher maximum cluster sum than the real data.

**Supplementary Figure 3:**
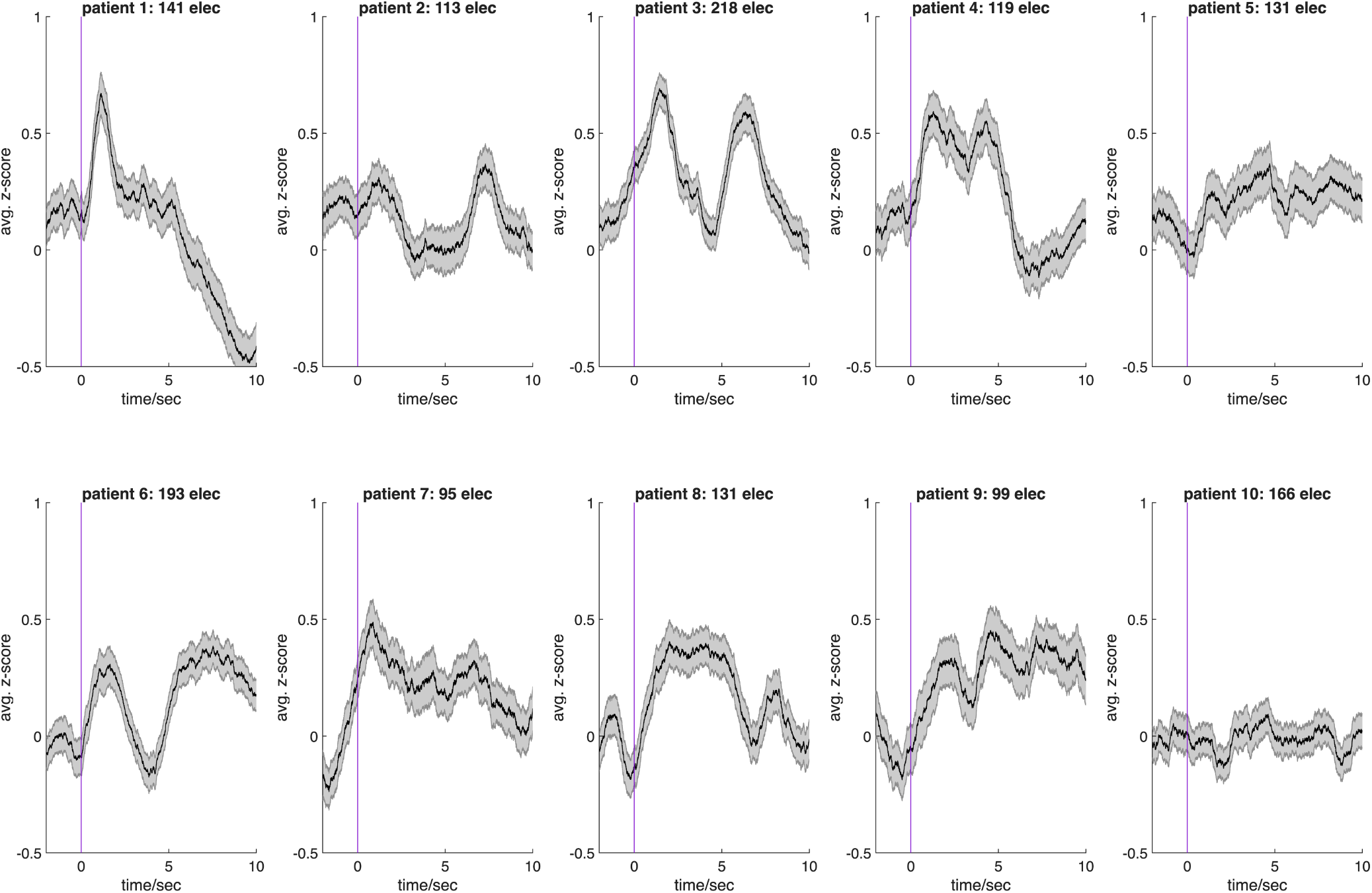
Patients’ individual time courses of z-scores measuring the association between neural state boundaries and the agreement on event boundaries in the norming sample.

**Supplementary Figure 4:**
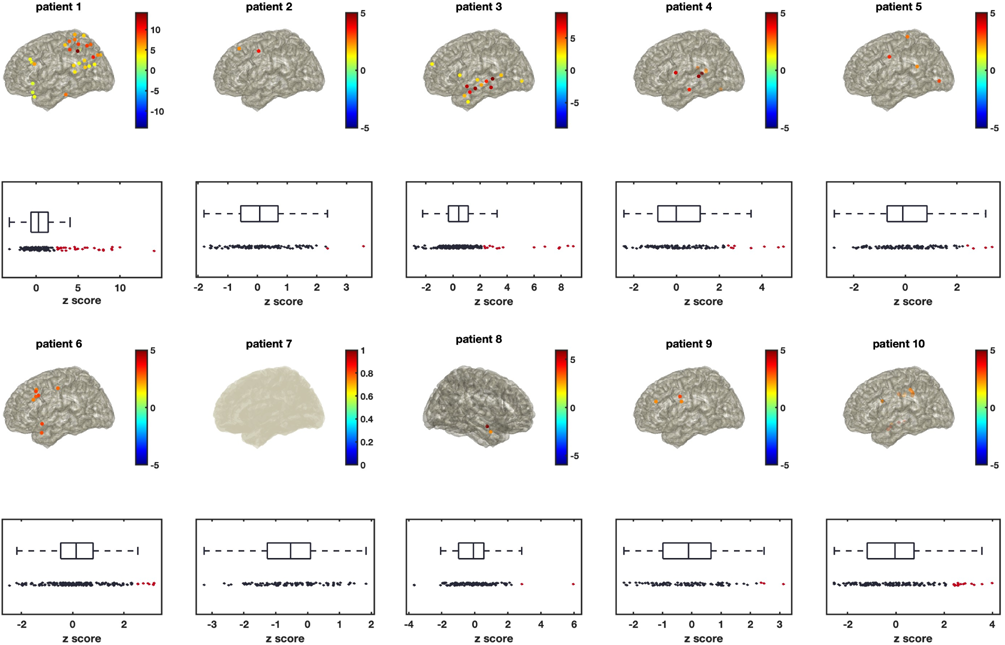
Patients’ individual CR-channels. Displayed is the z-score of reinstatement of state-patterns during the naturalistic interview superimposed on the brain (top) and displayed in red with box-plots of all channels (bottom).

## Footnotes

1 https://github.com/s-michelmann/semi_automatic_aligner

